# Genome-wide analysis of genetic risk factors for rheumatic heart disease in Aboriginal Australians provides support for pathogenic molecular mimicry

**DOI:** 10.1101/188334

**Authors:** Lesley-Ann Gray, Heather A D’Antoine, Steven Y. C. Tong, Melita McKinnon, Dawn Bessarab, Ngiare Brown, Bo Reményi, Andrew Steer, Genevieve Syn, Jenefer M Blackwell, Michael Inouye, Jonathan R Carapetis

## Abstract

**Background:** Rheumatic heart disease (RHD) following Group A Streptococcus (GAS) infections is heritable and prevalent in Indigenous populations. Molecular mimicry between human and GAS proteins triggers pro-inflammatory cardiac valve-reactive T-cells.

**Methods:** Genome-wide genetic analysis was undertaken in 1263 Aboriginal Australians (398 RHD cases; 865 controls). Single nucleotide polymorphisms (SNPs) were genotyped using Illumina HumanCoreExome BeadChips. Direct typing and imputation was used to fine-map the human leukocyte antigen (HLA) region. Epitope binding affinities were mapped for human cross-reactive GAS proteins, including M5 and M6.

**Results:** The strongest genetic association was intronic to HLA-DQA1 (rs9272622; P=1.86x10^−7^). Conditional analyses showed rs9272622 and/or DQA1*AA16 account for the HLA signal. HLA-DQA1*0101_DQB1*0503 (OR 1.44, 95%CI 1.09-1.90, P=9.56x10^−3^) and HLA-DQA1*0103_DQB1*0601 (OR 1.27, 95%CI 1.07-1.52, P=7.15x10^−3^) were risk haplotypes; HLA_DQA1*0301-DQB1*0402 (OR 0.30, 95%CI 0.14-0.65, P=2.36x10^−3^) was protective. Human myosin cross-reactive N-terminal and B repeat epitopes of GAS M5/M6 bind with higher affinity to DQA1/DQB1 alpha/beta dimers for the two risk haplotypes than the protective haplotype.

**Conclusions:** Variation at HLA_DQA1-DQB1 is the major genetic risk factor for RHD in Aboriginal Australians studied here. Cross-reactive epitopes bind with higher affinity to alpha/beta dimers formed by risk haplotypes, supporting molecular mimicry as the key mechanism of RHD pathogenesis.

## INTRODUCTION

Acute rheumatic fever (ARF) results from an autoimmune response to infections due to group *A Streptococcus* (GAS), *Streptococcus pyogenes*. Recurrences of ARF and its associated cardiac valvular inflammation lead to chronic valvular damage and rheumatic heart disease (RHD). RHD causes an estimated 275,000 deaths annually with an estimated 33 million prevalent cases globally (reviewed [1]). In Australia, RHD is most prevalent in the Indigenous population, affecting 2-6 per 1000 individuals (and as high as 15/1000 school aged children in the northern tropical regions[2])[3, 4].

The precise pathological mechanisms underlying RHD remain unclear. One hypothesis to explain inflammation of valvular tissue is molecular mimicry (reviewed [1, 5-8]). Accordingly, peptides from GAS proteins are processed by antigen presenting cells in the throat and heart tissue and presented on Human Leukocyte Antigen (HLA) class II molecules to CD4+ T lymphocytes that elicit pro-inflammatory cytokine responses and/or provide help to B lymphocytes for antibody secretion. In RHD patients, the CD4 T cell epitopes and antigenic specificities of antibodies show cross-reactivity to proteins in heart tissue, specifically targeting cardiac valves [5, 9]. This cross-reactivity is thought to be due to sequence similarities between heart tissues and GAS proteins, amongst which GAS M-proteins feature prominently [10], which is supported by studies of HLA-DQ-restricted T cell clones that recognise the M protein and myosin peptides in the blood and hearts of RHD patients [11, 12] as well as studies in animal models of disease [13]. The precise mechanism by which these cross-reactive antibodies target the valve is unclear, and cross-reactive antibodies have been observed in streptococcal pharyngitis without complications [14]. An alternative hypothesis (reviewed [7]) is that a streptococcal M protein N-terminus domain binds to the CB3 region in collagen type IV, initiating an antibody response to the collagen which results in inflammation. These antibodies do not cross-react with M proteins, and hence do not involve molecular mimicry.

Key aspects of molecular mimicry are the relevant proteins/peptides in GAS strains and host susceptibility. In the Northern Territory of Australia there is high genetic diversity amongst GAS strains which reflect global-scale transmission rather than localised diversification [15, 16]. Despite ubiquitous exposure to GAS, only 1-2% of Indigenous Australians living in this region develop RHD, and the cumulative incidence of ARF only reaches 5-6% in communities with the most complete case ascertainment [17]. ARF is a precursor to RHD, and in a meta-analysis of 435 twin pairs susceptibility to rheumatic fever was estimated to be 60% heritable [18]. For RHD, a number of candidate gene studies have variably reported associations with genes controlling innate and adaptive immune responses (reviewed [6]). Among these candidates, HLA Class I and II genes feature most prominently, but with little consistency in risk and protective genes/alleles reported [6, 19, 20]. Recently, a GWAS of RHD was performed in Oceania populations but did not report an HLA signal [21]. This variability in reported associations likely reflects differing study designs, population-related genetic heterogeneity, failure to control for confounding factors, and the vagaries of small samples sizes and candidate gene approaches. Here we undertake an unbiased genome-wide approach to identify genetic risk factors for RHD in echocardiogram-confirmed cases from the Northern Territory of Australia. The HLA-DQA1-DQB1 locus was the only region to show strong association in this population. We show that differential binding of GAS/human cross-reactive epitopes to MHC Class II dimers for specific HLA DQA1_DQB1 risk and protective haplotypes may underpin the molecular mimicry hypothesis for RHD pathogenesis.

## METHODS

### Ethical Considerations, Sampling and Clinical Data Collection

This study was undertaken with ethical approval from the Human Research Ethics Committee (HREC) of the Northern Territory Department of Health and Menzies School of Health Research (ID HREC-2010-1484) and the Central Australian HREC (ID HREC-2014-241). The study was overseen by a project steering committee and three sub-committees – Aboriginal governance, clinical and scientific. The protocol and any key changes required agreement from the Aboriginal governance committee. Stage 1 of the project involved community engagement and consent, development of culturally appropriate consent materials, and establishment of appropriate governance for collection and subsequent storage of samples. Stage 2 involved identifying individual participants, obtaining informed consent, and collection of samples and associated meta-data. The individual consent incorporated an “opt-in” design where participants selected which components of the study they were comfortable to participate in, and were able to withdraw from the study at any stage [22]. This included an option to accept or refuse continued use of their genetic or clinical data in further studies. De-identified post QC (cf. below) genotype data for individuals who consented to continued use of their data have been lodged in the European Genome-phenome Archive (accession number EGAS0000000000) with access controlled through a study-specific Data Access Committee.

Participants were recruited from 19 communities in the Northern Territory of Australia (Figure 1). Case participants were defined a priori as having had, at some stage, echocardiographically confirmed evidence of RHD and/or ARF with carditis. For each of the 19 communities, we obtained a list of individuals on the Northern Territory Rheumatic Heart Disease register. These lists were further screened for patients with a history of ARF and associated carditis (defined using the 2015 revised Jones Criteria [23]) or RHD confirmed on echocardiogram (defined using the 2012 World Heart Federation criteria [24]). We aimed for a 1:2 ratio of cases to controls. Controls were selected from the same communities (range 4-215 participants/community) to ensure similar likelihood of exposure to GAS among cases and controls, and included a selection of family members as well as unrelated community-based controls. Medical records of potential control participants were checked to exclude a prior history of rheumatic fever. We did not perform echocardiograms on control participants. Both cases and controls had to be aged ≥18 years, to minimise the likelihood of enrolling controls that might subsequently become cases (given that ARF is largely a disease of school aged children and most RHD cases are diagnosed before the age of 30). Data were collected for age, gender, community location and RHD case/control status.

**Figure 1.**
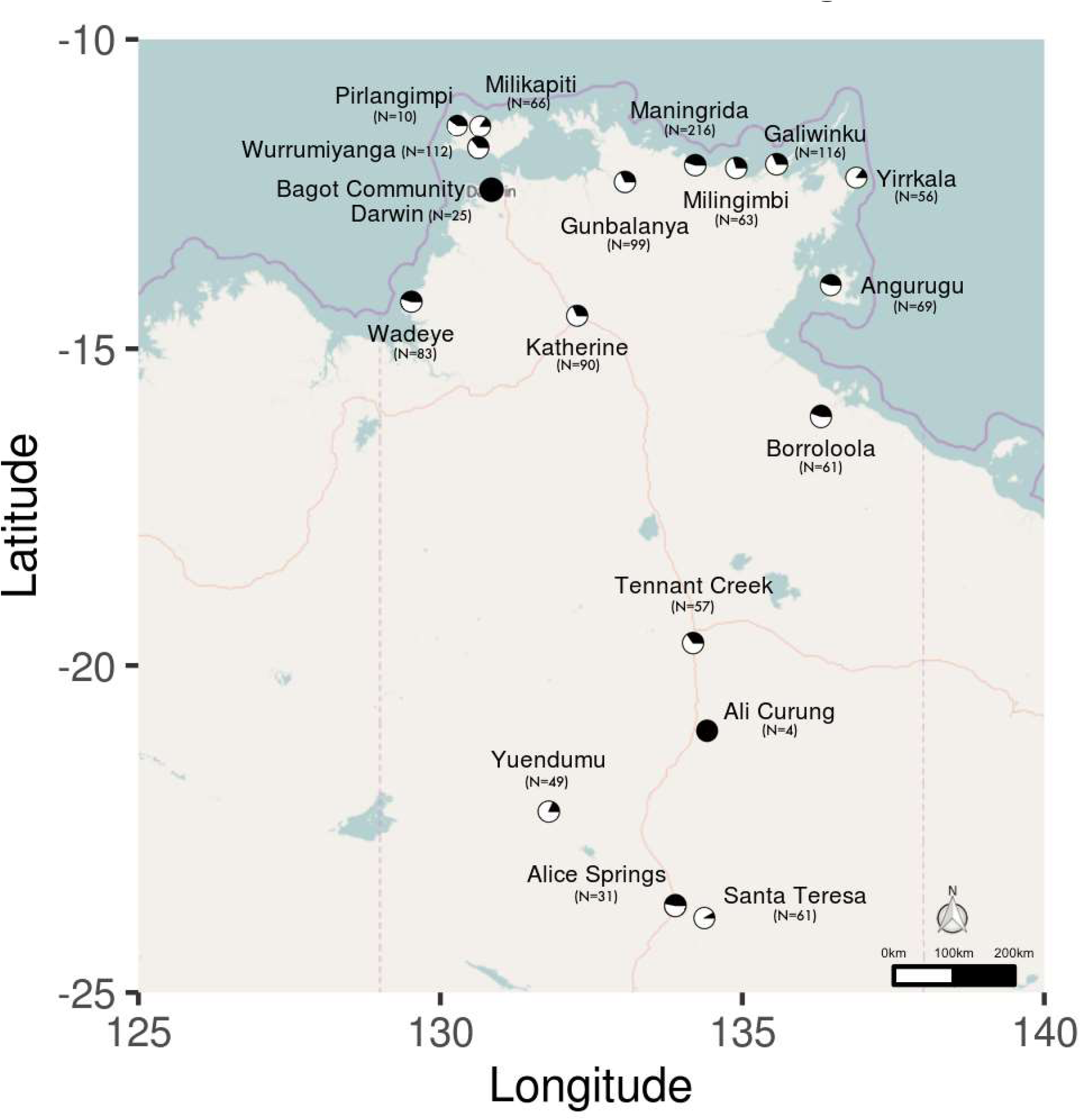
Locations of study populations. Locations are given by latitude and longitude for 19 Aboriginal communities in the Northern Territory of Australia that participated in the study. Each dot indicates a single community, with wedges indicating the proportion of case (filled in wedge) compared to control (open wedges) samples for each population.

We collected clinical data and saliva from 1382 individuals. Of these, 11 later withdrew consent for the study, and an additional 71 individuals were deemed ineligible for case or control status following detailed medical record review, leaving 1291 eligible to include in the study. Demographic details (age, sex, case/control status) for the 1263 (of 1291) study participants who also passed QC following genotyping (cf. below) are summarised in Table S1.

### Array Genotyping and Marker QC

Saliva was collected using Oragene OG-500 saliva kits (DNA Genotek Inc., Ontario, Canada) and DNA extracted according to manufacturer protocols. DNAs were genotyped on the Illumina Infinium^®^ HumanCoreExome Beadchip (Illumina Inc., San Diego, CA, USA), which includes probes for 547,644 single nucleotide polymorphisms (SNPs), 281,725 of which are genome-wide tag SNPs that represent core content and are highly informative across ancestries, and 265,919 SNPs that are exome-focused markers. All genotyping data and reference panels were analysed using human genome build 37 (hg19). Individuals were excluded if they had a missing data rate >5%. SNP variants were excluded if they had genotype missingness >5%, minor allele frequency (MAF) <0.01, or if they deviated from Hardy-Weinberg equilibrium (HWE; threshold of P<1.0x10^-6^). This provided a post-QC dataset of 1263 individuals genotyped for 239,536 markers. This sample comprised 398 cases and 865 controls (Table S1), providing 68% power to detect genome wide significance (P<5x10^-8^) for genetic effects with a disease allele frequency of 0.25, effect size (genotype relative risk) of 2, and assuming a disease prevalence of 2%. Overt non-Aboriginal population stratification was assessed using the top 10 principal components (PCs) from FlashPCA [25].

### SNP Imputation and GWAS

Imputation of missing and unassayed genetic variants was performed using the 1000 Genomes Project phase 3 reference panel [26], which contains 88 million variants for 2502 samples from 26 populations throughout Africa, America, East Asia, Europe and South-East Asia. Array variants were phased using SHAPEIT v2 (r644) [27] and imputed with IMPUTE v2.3.2 [28]. We excluded imputed SNPs with an information metric <0.4 or genotype probability <0.9, and the remaining variants were converted to genotype calls and filtered for <10% missingness and MAF>0.01. Imputation accuracy was assessed using the r^2^ metric (r^2^>0.8), which represents the squared Pearson correlation between the imputed SNP dosage and the known allele dosage.

Genome-wide association analysis for the RHD phenotype was performed using a linear mixed model as implemented in FaST-LMM v2.07, which takes account of both relatedness and population substructure [29]. Age and gender were included as fixed effects in the model. Population structure and relatedness were controlled using the genetic similarity matrix, computed from 41,926 LD-pruned array variants, and any systematic confounding assessed using QQ plots and a test statistic inflation factor (λ). Genome-wide significance was set at P<5x10^-^8 [30].

### Fine-Mapping Associations in the HLA Region

Conditional association analyses in the HLA region also utilised FaST-LMM. Univariate conditional analysis can fail to uncover residual signals due to the long-distance haplotypes observed in the HLA region [31], therefore we used a step-wise conditional analysis of classical HLA alleles and amino acids to scan for independent signals in HLA. First, we typed exons of 10 classical HLA alleles for 716 samples using the TruSight HLA sequencing panel and produced 4-digit phase-resolved genotype calls against the IMGT v3210 database (Murdoch University Centre for Clinical Immunology and Biomedical Statistics, Perth, Western Australia). We generated an Aboriginal reference panel of typed HLA variants from these individuals and imputed the HLA region for the untyped individuals using HIBAG [32]. Phased genotype calls with *prob* >0.8 (i.e. conditional probability of pairs of haplotypes consistent with observed genotypes) were converted to amino acid variants and merged with the SNP variants for association analysis in FaST-LMM, as described above. Haplotype analyses were performed in PLINK [33] on phased haplotype data using logistic regression under an additive model with gender, age and 10 PCs as covariates.

### Functional Predictions for Candidate Loci

We assessed the functional role for the candidate causal HLA variants *in silico* using NetMHCIIPan 3.1 [34] to map epitopes and their binding affinities to two risk and one protective HLA-DQA1_HLA-DQB1 haplotypes across GAS proteins known to contain human cross-reactive epitopes. A literature review of the GAS proteins reported to show cross-reactivity with host tissue proteins was undertaken (Table S2). Full-length amino acid sequences of all GAS proteins, including M5 and M6 proteins, shown to have cross-reactive epitopes were converted to a series of 20-mer sequences with a 1-mer sliding window and assessed for binding to each significantly associated DQA1_DQB1 haplotype. Cross-reactive epitopes from human proteins were mapped onto the epitope binding maps of M5 and M6, as indicated. Binding affinities were compared (GraphPad Prism 7.00: 1-way ANOVA with multiple comparisons and correction for multiple testing) between haplotypes across the regions of peak epitope binding where 20-mer epitopes shared common 9-mer core epitopes.

## RESULTS

### Genome-wide Association Study

We conducted a GWAS for rheumatic heart disease (RHD) in 1263 individuals comprising 398 RHD case and 865 control participants from communities in the Northern Territory of Australia. From direct genotyping on the Illumina HumanCoreExome array, we achieved 4.46 million high quality imputed variants (92.33% of variants imputed to high accuracy, r^2^>0.80) with moderate to high imputation accuracy genome-wide (Figure S1A). Genetic population structure was clearly evident from principal components analysis, largely capturing the geographic distribution of the remote Aboriginal Australian communities (data not shown). The use of a linear mixed model framework with genetic relatedness matrix (FastLMM) to perform a genome-wide association study for RHD effectively controlled this stratification, as evidenced by a quantile-quantile plot of the p-values from the genome-wide scan (λ =1.021; Figure S1B). A single major signal was detected within the class II region of the HLA gene family on chromosome 6 which peaked at the imputed variant rs9272622 (32607986bp, *P*=1.86x10^−7^, OR=0.897 for protective allele C) within intron 1 of *HLA-DQA1* (Figure 2).

**Figure 2.**
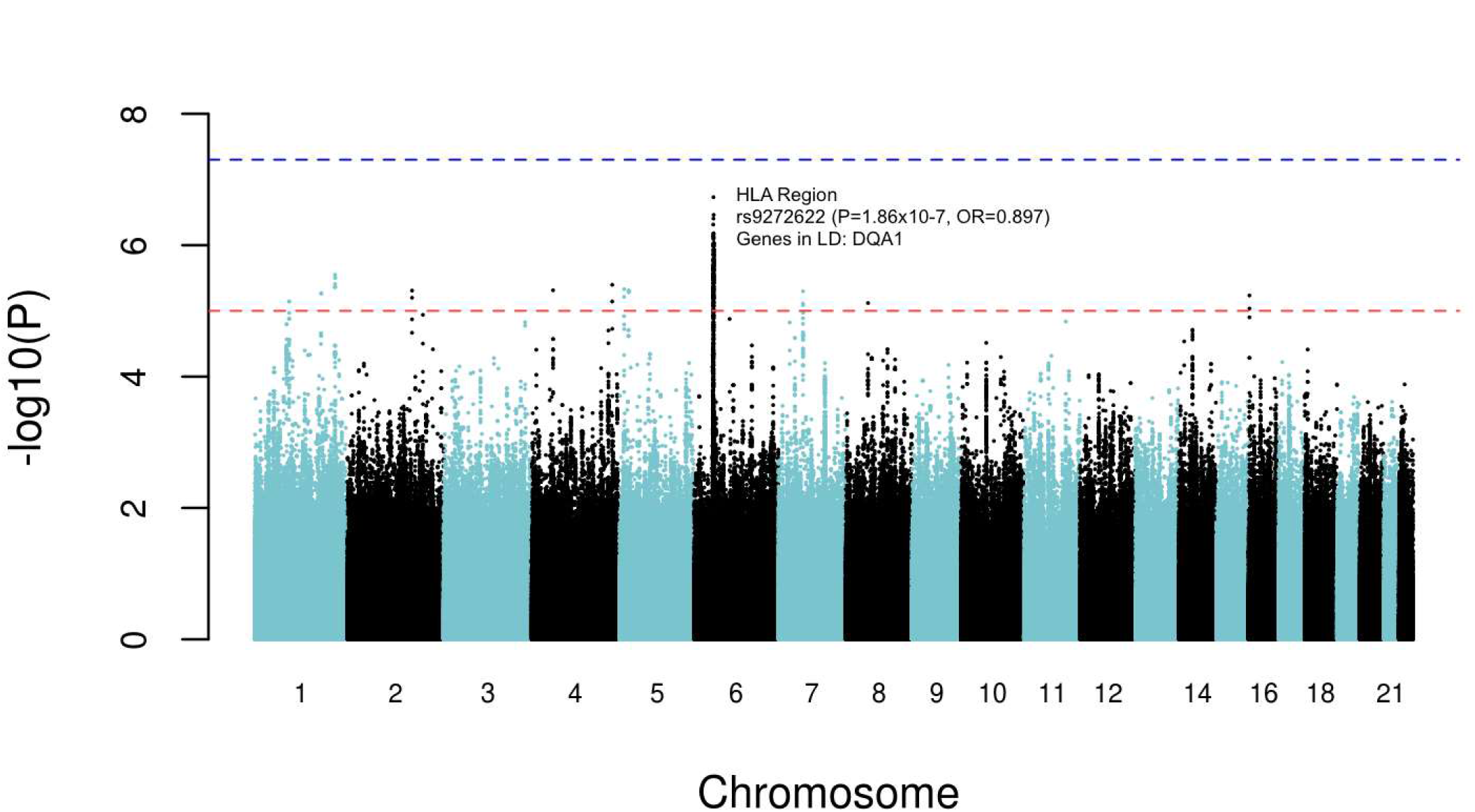
Manhattan plot of GWAS results for the 4.46M high quality 1000G imputed SNP variants. Data are for analysis in FastLMM looking for association between SNPs and RHD. The top SNP rs9272622 occurred within the HLA region on Chromosome 6p21, as shown.

### Fine-mapping the HLA Class II Region

Regional plots of the class II region showed that the top SNP rs9272622 tagged a linkage disequilibrium block (r^2^>0.8) across the *HLA-DQA1* to *HLA-DQB1* region (Figure 3). There were no residual signals across the HLA Class II region after conditioning on the index variant rs9272622 (Figure S2). To further understand the potential functional variants across the HLA Class II region, we typed and imputed traditional 4-digit HLA alleles, converted alleles to amino acid calls, and applied a multiple stepwise regression analysis. The top 4-digit HLA alleles for risk and protection were *HLA-DQB1*0601* (*P*=4.06x10^−4^, OR=1.07) and *HLA-DQA1*0301* (*P*=2.71x10^−4^, OR=0.92), respectively. The top 4-digit HLA-DRB1 association was HLA-DRB1*0803 (*P*=0.005, OR=1.06), and no significant associations were observed for classical alleles across the SNP poor region (Figure 3) of HLA-DRB3/DRB4/DRB5. The strongest amino acid associations (Figure 4A) were at positions AA_DQA1_16_32713236 (*P*=2.08x10^−6^, OR=0.91) and AA_DQA1_69_32717257_L (*P*=2.08x10^−6^, OR=0.91) in exons 1 and 2 of DQA1, respectively, which were in 100% linkage disequilibrium with each other, and at AA_DQB1_38_32740723 (*P*=2.17x10^−6^, OR=0.91) in exon 2 of DQB1. As when conditioning on the top SNP (Figure 4B), there was no residual signal across the HLA region when conditioning on either the top DQA1 AA variant (Figure 4C) or both the top SNP and the top DQA1 AA variant (Figure 4D), suggesting that associations across the HLA-DQA1 to HLA-DQB1 region are all due to linkage disequilibrium with top variants at *HLA-DQA1*.

**Figure 3.**
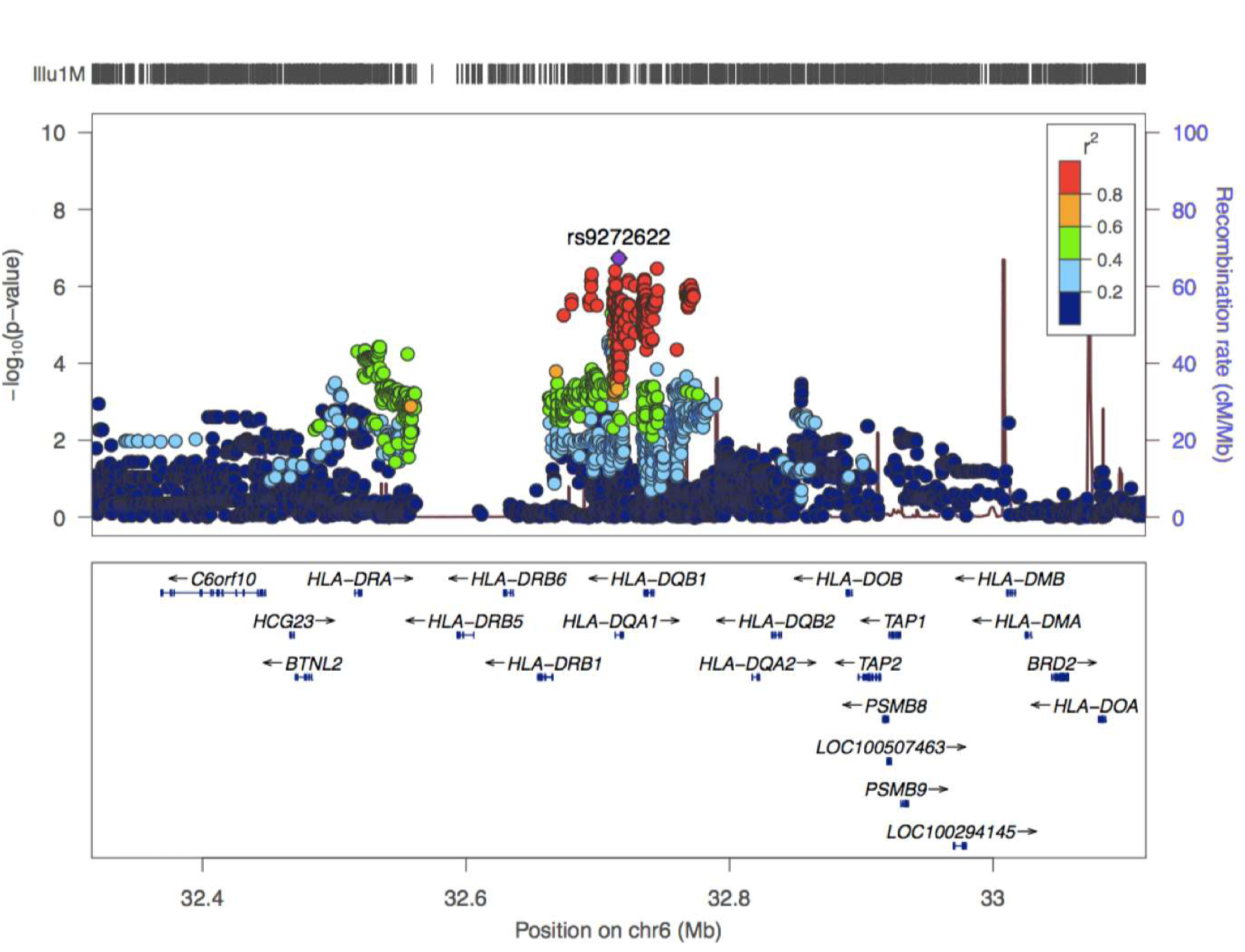
LocusZoom plot of SNP associations with RHD across the Class II region of the HLA complex. The–log_10_P values (left Y-axis) are shown in the upper section of the plot. Dots representing individual SNPs are coloured (see key) based on their linkage disequilibrium r^2^ with the top SNP rs9272622. The right Y-axis is for recombination rate (blue line), based on HapMap data. The bottom section of the plot shows the positions of genes across the region. For clarity, 5 genes were removed upstream of 32.8Mb (PSMB8-9, HLA-DOA, LOC100507463, LOC100294145).

**Figure 4.**
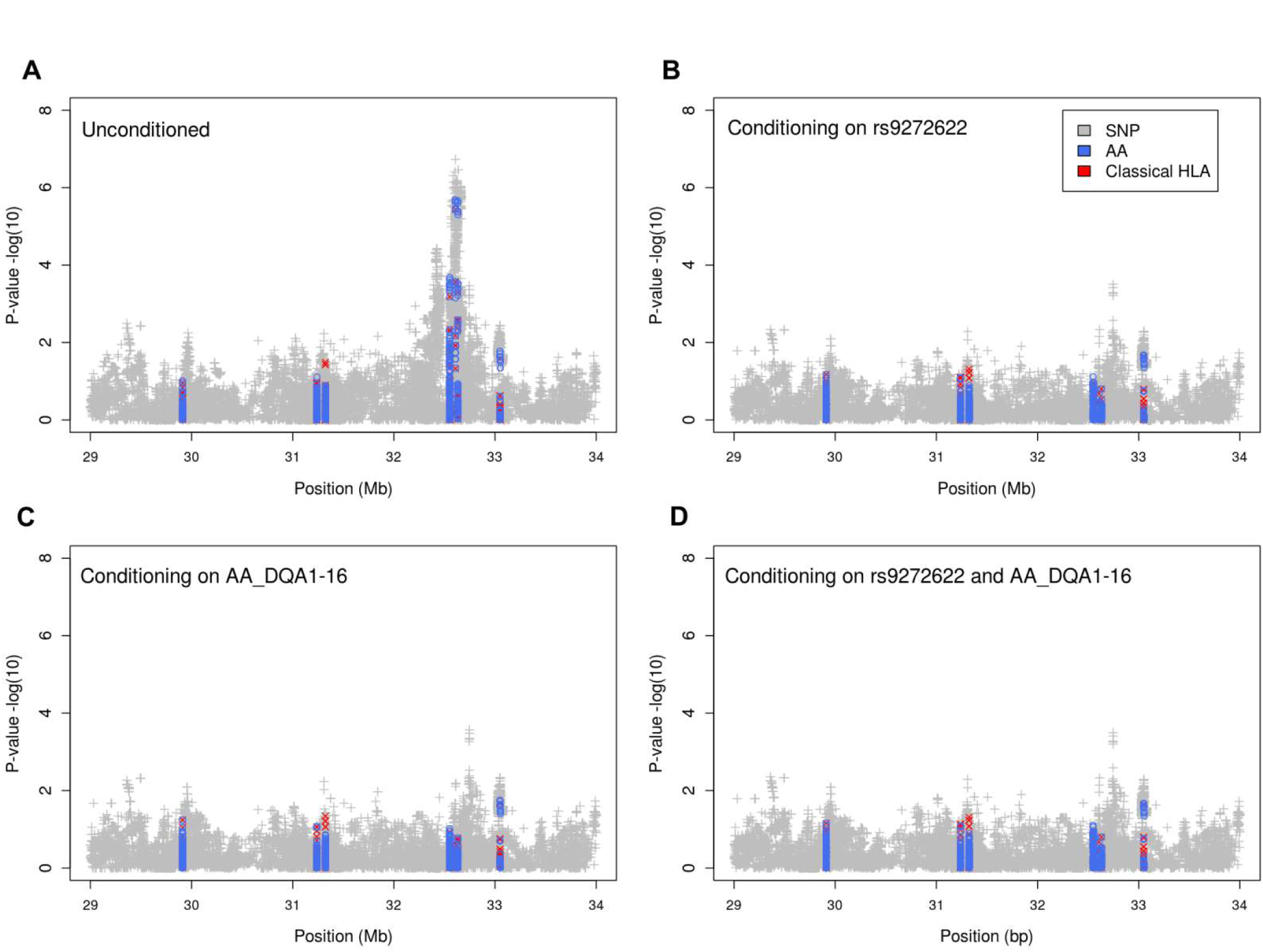
Plots of association between RHD and imputed classical 4-digit and amino acid (AA) HLA alleles. Results are for association analyses in FastLMM: (A) without conditioning; (B) after conditioning on the top SNP rs9272622; (C) after conditioning on the top AA variant at DQA1 AA position 16; and (D) after conditioning on both of these variants.

### HLA-DQ haplotype risk

*HLA-DQA1* and *HLA-DQB1* genes encode alpha and beta chains, respectively, forming DQ alpha/beta heterodimers that together bind antigenic epitopes to present to CD4+ T cells. For antigen presentation via HLA-DQ class II molecules, variation at both the alpha and beta chains contribute to epitope binding to the peptide groove encoded by exons 2 of both alpha and beta chains. Variants at both genes may thus contribute together to determine risk versus protection from RHD. We therefore looked for associations between RHD and *HLA-DQA1_HLA-DQB1* haplotypes. Haplotype analysis in PLINK identified HLA-DQA1*0101_DQB1*0503 (OR 1.44, 95%CI 1.09-1.90, P=9.56x10^−3^) and HLA-DQA1*0103_DQB1*0601 (OR 1.27, 95%CI 1.07-1.52, P=7.15x10^−3^) as risk haplotypes, with HLA-DQA1*0301_DQB1*0402 (OR 0.30, 95%CI 0.14-0.65, P=2.36x10^−3^) as the protective haplotype for RHD in the study population (Figure 5). These haplotypes were taken forward in *in silico* functional analyses.

**Figure 5.**
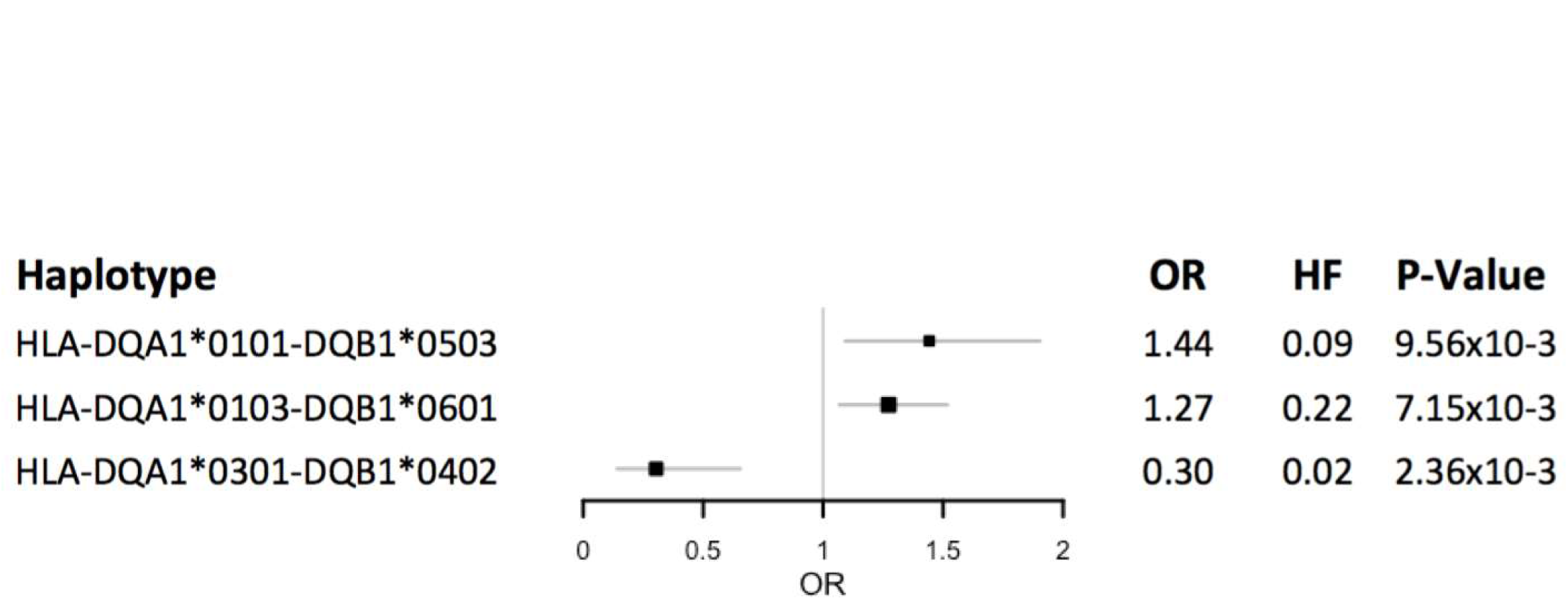
Forest plot showing associations between RHD and phased HLA DQ_DB haplotypes. The plot show odds ratios (OR) and 95% confidence intervals for two risk (OR>1) and one protective (OR<1) haplotypes. Information to the right of the plot shows values for the OR, the haplotype frequency (HF), and the p-value for the haplotype association.

### Mapping Group A Streptococcus epitopes to risk versus protective HLA DQ haplotypes

There are two important ways in which association between HLA-DQ haplotypes could impact on disease susceptibility and control programs: (i) in the pathogenesis of disease, particularly in relation to an autoimmune mechanism for RHD through GAS epitopes that cross-react with self; and (ii) in the ability of high risk individuals to respond to proposed vaccine antigens. To address the first, we initially assessed the binding affinities of epitopes across the M-proteins M5 and M6 from rheumatogenic GAS strains to the alpha/beta heterodimers specific to the observed risk versus protective HLA-DQA-HLA-DQB haplotypes. Figure 6 shows the epitope binding affinities mapped for these haplotypes across the full-length M5 and M6 proteins, together with annotation indicating the positions along each protein where experimentally validated cross-reactive epitopes have been identified (Table S2). Several epitope peaks that correspond to key cross-reactive epitopes are shown to bind with higher affinity to the two risk haplotypes compared to the protective haplotype (Figure 6), notably in the B repeat regions previously shown to contain key cross-reactive T cell epitopes with human cardiac myosin (e.g. Cunningham et al., 1997 [10]; see also Table S2). The peak differences in binding affinities for 20-mer epitopes in these regions of previously experimentally validated cross reactivity for the M5 (see arrows, Figure 6A) and M6 (see arrows, Figure 6B) proteins were highly significant (P<0.0001) between risk and protective haplotypes (Figure 7). No differences in epitope binding to risk versus protective haplotypes were observed when we mapped epitopes across GAS M proteins (e.g. E pattern M4 and M49 types [35]) from non-RHD GAS strains (Figure S3). Nor did we observe regions of differential epitope binding affinities across other GAS proteins (HSP70, STRP1; Figure S3) reported in the literature to contain epitopes cross reactive with human proteins implicated in RHD pathogenesis (Table S2).

**Figure 6.**
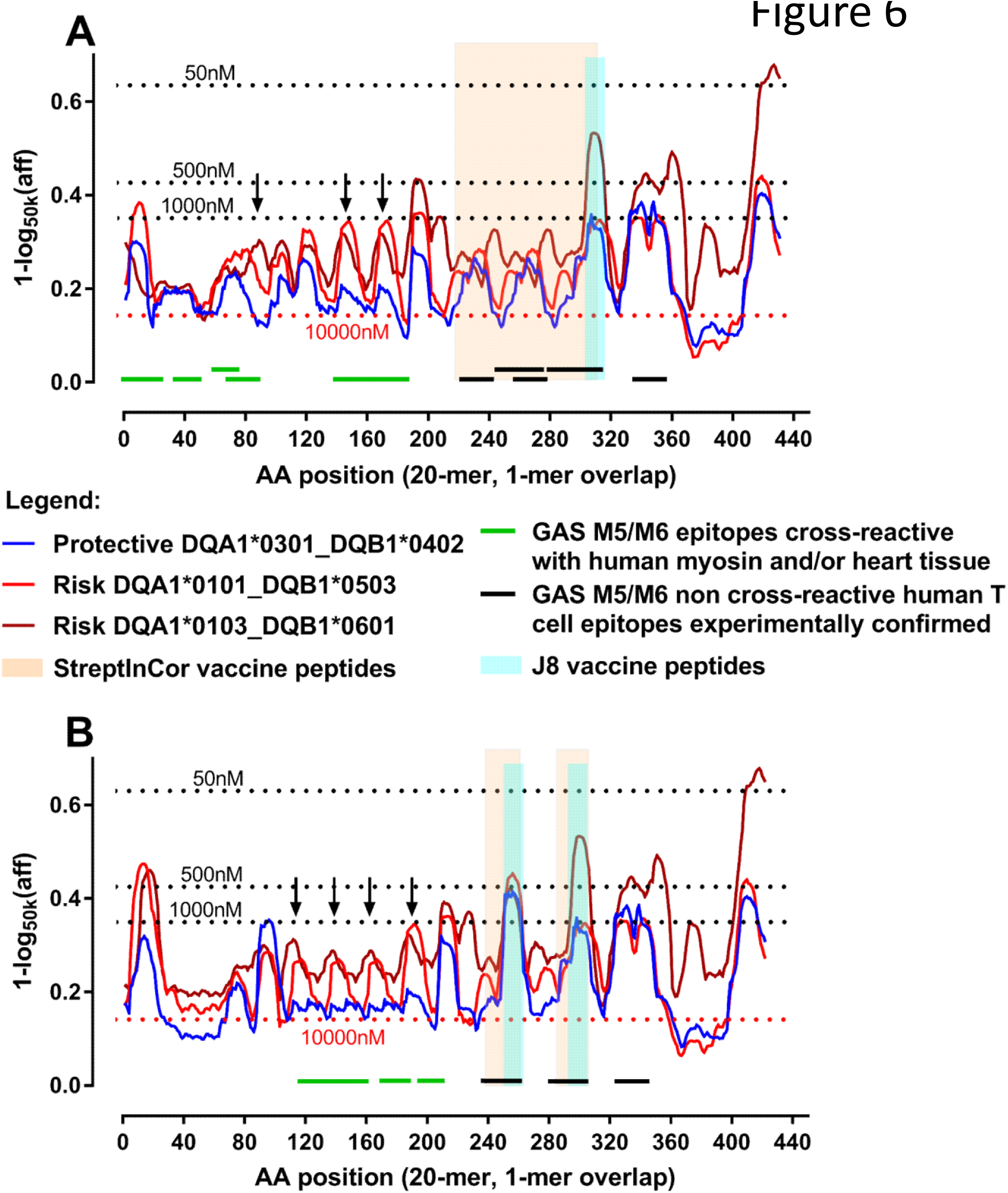
Plots showing binding affinities for predicted epitopes of GAS M proteins recognised by HLA DQ-DB heterodimers. Epitope binding predictions were performed in NetMHCIIPan3.1. The y-axis shows the relative binding affinity (expressed as 1-log_50,000_ of the nM binding affinity) for heterodimers formed from risk (red, brown) and protective (blue) DQ_DB haplotypes (see key); the x-axis indicates the amino acid sequence locations for mature proteins, also equivalent to the start position of overlapping 20mers (1-mer sliding window) in (A) the GAS M5 sequence and (B) the GAS M6 sequence. Horizontal dotted lines show different nM binding affinities. Negative binding affinity is indicated at >10,000 nM (i.e. below the red dotted line). Vertical arrows indicate the N-terminal or B-repeat cross-reactive epitopes used to compare binding affinities in Figure 7. The linear positions of known cross-reactive epitopes with human cardiac myosin and/or human heart valve tissue are shown in green; black lines indicate the regions of known experimentally determined human T cell epitopes (see Table S2). The apricot and pale blue vertical bars indicate the positions of C-repeat region peptides incorporated into the StreptinCor and J8-DT vaccines, respectively.

**Figure 7.**
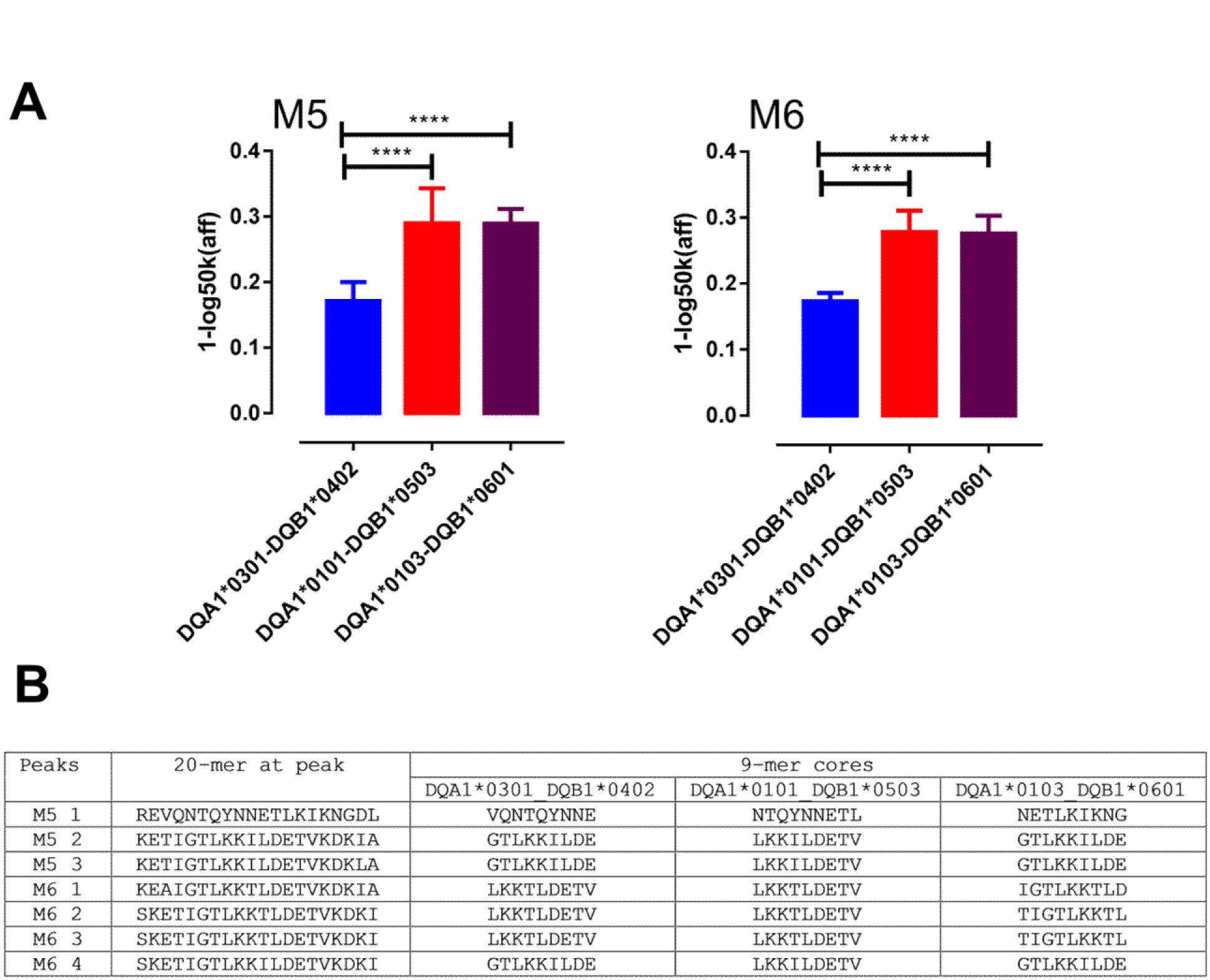
Mean binding affinities for GAS M protein epitopes cross-reactive with human cardiac myosin. (A) The y-axis (as for Figure 6) shows mean plus SD for predicted M5 and M6 GAS protein epitopes (NT and B repeat regions; as annotated with arrows in Figure 6) recognised by risk (red and brown bars) or protective (blue bar) DQ-DB heterodimers formed from DQA1_DQB1 haplotypes, as labelled. **** indicates P<0.0001. (B) Shows the 20-mer epitope at the peak of the differences for binding affinity of risk versus protective haplotypes, together with the predicted 9-mer cores for each haplotype.

Also annotated in Figure 6 are the C-terminal regions of the M5/M6 proteins that contain peptides incorporated into the two candidate vaccines currently in advanced stages of development that include antigens from this M protein region, J8-DT [36] (vertical blue strip) and StreptinCor [37] (vertical apricot strip). Whilst the risk haplotype HLA-DQA1*0103_DQB1*0601 binds to epitopes across this region with higher affinity, all three haplotypes show similar patterns of epitope binding across this region. None show the low level of binding affinity such as that observed for the protective haplotype for cross-reactive epitopes across the B-repeat region. These results suggest that individuals genetically at risk of developing RHD have the potential to make HLA-DQ-driven CD4+ T cell responses to these vaccines.

## DISCUSSION

The results of an unbiased genome-wide evaluation of genetic determinants for RHD in Aboriginal Australians living in northern Australia provide evidence for a prominent association in the class II gene region of HLA, consistent with prior data from more limited genetic studies. Strong linkage disequilibrium across HLA, together with variable selection of candidate HLA genes, likely contributes to the inconsistency in the HLA genes/alleles associated with risk versus protective from RHD in prior studies [6, 19, 20] even though experimental studies support HLA-DQ restriction of T cell clones involved in T cell mimicry in RHD [11]. In contrast, our study benefitted from dense fine mapping across HLA, allowing us to identify specific risk (HLA-DQA1*0101_DQB1*0503; HLA-DQA1*0103_DQB1*0601) versus protective (HLA-DQA1*0301_DQB1*0402) haplotypes across the genes encoding alpha and beta chains of HLA-DQ. While our conditional analysis suggested only a single HLA signal, we cannot discount the possibility that other genes may contribute to genetic susceptibility to RHD in this population. It is of specific interest, however, that our study did not find evidence for replication for variants at the IGH locus recently shown to be significantly associated with RHD in a GWAS of New Caledonian and Fijian populations [21]. Differences in study design and phenotype classification may have contributed, as could genetic heterogeneity between indigenous populations which is known to occur for autoimmune and infectious diseases [38]. It is reassuring, nevertheless, that both GWAS have found evidence consistent with autoimmune genetic architecture. Ultimately, meta-analyses of greater statistical power will be required to investigate population-specific differences and detect additional RHD loci.

Our identification of risk versus protective haplotypes across HLA-DQA/DQB provided an opportunity to revisit the molecular mimicry hypothesis in relation to RHD pathogenesis. Dimers created from alpha and beta chains of HLA class II molecules present epitopes processed from foreign proteins to CD4 T cells, the preferred outcome of which would be to provide an immune response that will protect against infection. In the context of autoimmune disease, self-epitopes are presented and recognised as non-self, leading to detrimental immune pathology. The molecular mimicry hypothesis proposes that GAS contains proteins with AA sequences that mimic (or are cross-reactive with) human proteins, thus leading the immune system to recognise them as auto-antigens that drive immune pathology rather than (or in addition to) immunity against GAS itself.[1, 6] In the case of HLA-DQ, variation in exons 2 of both alpha and beta chains encoded by DQA and DQB, respectively, contribute to variation in shape and structure of the epitope binding pocket [39]. This means that the specific alpha/beta dimers encoded by DQA/DQB genes carried on the same haplotype will create binding pockets that have different characteristics in terms of ability to bind and present epitopes to CD4+ T cells. Using the current gold standard NetMHCIIPan 3.1[34, 40] predictive algorithm to map specific epitopes across GAS proteins allowed us to identify significant differences in the ability of dimers created from risk versus protective haplotypes to bind cross-reactive epitopes. In particular, cross-reactive epitopes from cardiac myosin, one of the key cardiac proteins thought to contribute to the molecular mimicry hypothesis in RHD [1, 6, 10], were predicted to bind to dimers created from risk haplotypes but have no predicted binding to dimers created from the protective haplotype. Thus we identify a potential molecular mechanism to account for immune pathogenesis causing RHD in this population. Although we carried out our epitope mapping studies on just two M5 and M6 GAS strains most studied for the presence of human cross-reactive epitopes, our results are relevant to all GAS strains carrying cross-reactive N-terminal or B repeats. Relevance to our study population is consistent with global-scale transmission of GAS strains in this remote Aboriginal population [15]. Of interest too is the observation that, whilst rare cases of dimers created by *trans* association of alpha/beta chains encoded on opposite strands of the chromosome have been observed to contribute to susceptibility to type 1 diabetes, the predominant observation is that dimers are formed by alpha/beta chains encoded in *cis* [39]. This likely contributes to our ability to identify risk versus protective haplotypes across the HLA DQA1-DQB1 region, since strong linkage disequilibrium will keep particular combinations of DQA/DQB genes together in *cis*.

More broadly, this study represents a rare example of a genome-wide association study in a remote Indigenous population, yet one which shows that such studies can be successfully undertaken and uncover insights which have the potential to inform pathogenesis and vaccination strategy.

In conclusion, we here present results of the first GWAS undertaken for RHD in an Aboriginal Australian population. We report strong evidence for a role for HLA DQ/DB Class II molecules, and we link this to significant differences in affinity of binding of cross-reactive epitopes from GAS M proteins to antigen presenting heterodimers formed by risk versus protective DQ-DB haplotypes. Further functional analysis of T cell responses to cross-reactive T cell epitopes, as carried out in previous studies [11], could now be targeted at these specific DQ-DB heterodimers. Overall, our results provide new data on mechanisms that may contribute to risk of RHD caused by GAS strains.

## Author contributions

L-AG, HD’A, and SYCT contribute equally to the work. JMB, MlI, and JRC contributed equally to supervision of the work. L-AG managed the data and carried out the genetic statistical and bioinformatic analyses, and prepared the first draft of the manuscript. HD’A and MMcK project managed in Darwin, including management of ethical, legal and social aspects of the study. MMcK carried out the field work and sample collection. DB and NB made significant contributions to governance and helped design the community engagement arms of the project. SYCT, BR and AS provided the major clinical inputs for diagnosis and review of patient records. GS prepared the DNAs including quality control, liaised with providers for both chip genotyping and sequence-based HLA typing. JMB and MlI supervised the GWAS and HLA fine mapping analysis. JMB devised, supervised and interpreted the *in silico* analyses, and undertook major revisions of manuscript. JRC was the lead investigator on the project. All authors reviewed and approved the final manuscript.

## Acknowledgements

We would like to acknowledge all Chief Investigators of this study, the project team including the community based researchers, the communities, agencies and all the participants for their invaluable contribution to this project. We also acknowledge the contribution of Paul I.W. de Bakker, now Vertex Pharmaceuticals, to the design and initial analysis of the study, and thank Kara Imbrogno and Grace Chua for assistance with preparation of some of the DNAs used for this study.

---

**Figure S1.**
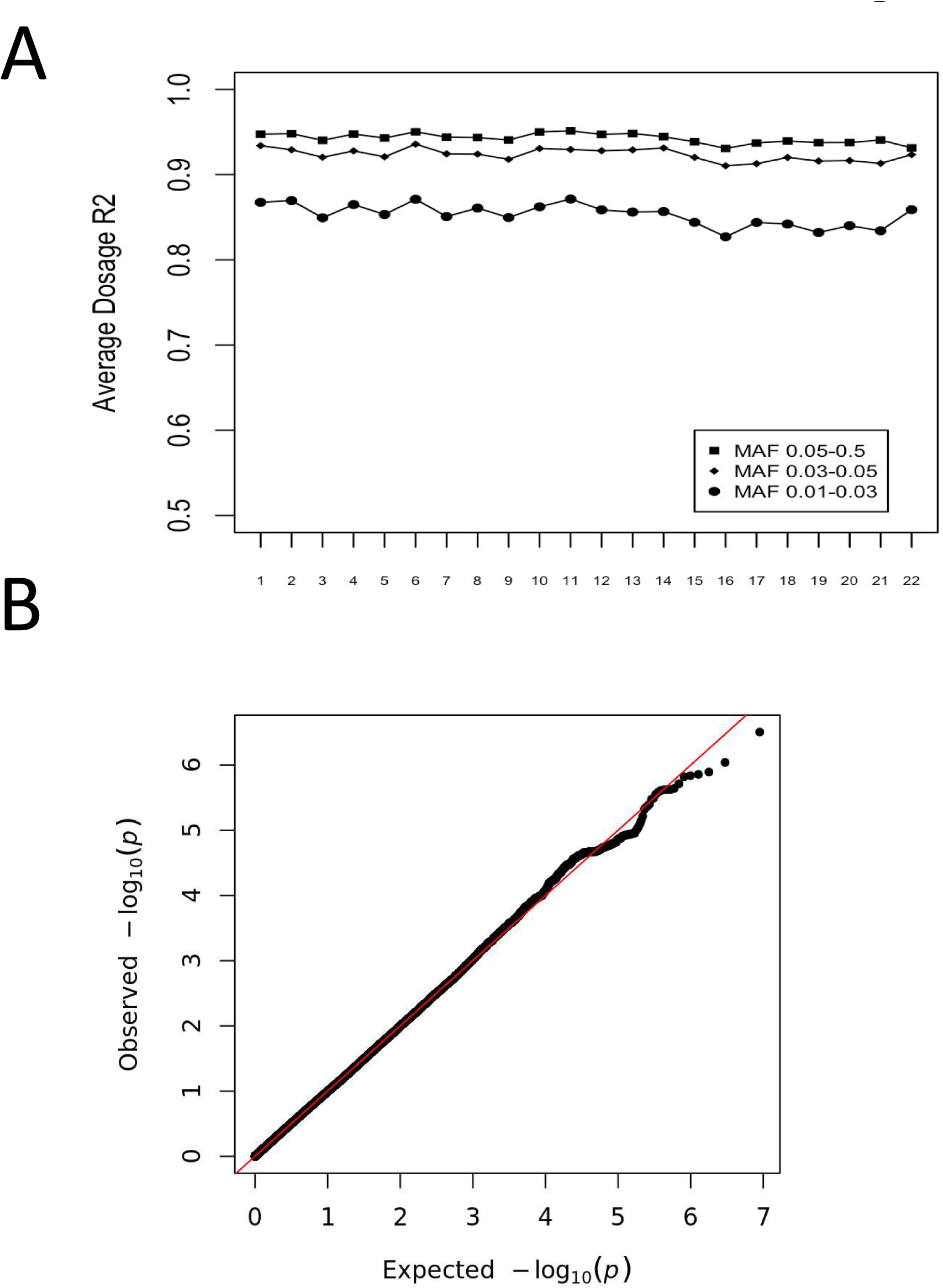
(A) Imputation accuracy measured as average dosage R^2^ for 235,942 type 2 variants filtered for an information metric >0.4 and genotype probability call > 0.90. Data shown separately by chromosome and for different minor allele frequency (MAF) ranges, as indicated in the key. (B) Quantile-quantile plot of GWAS p-values.

**Figure S2.**
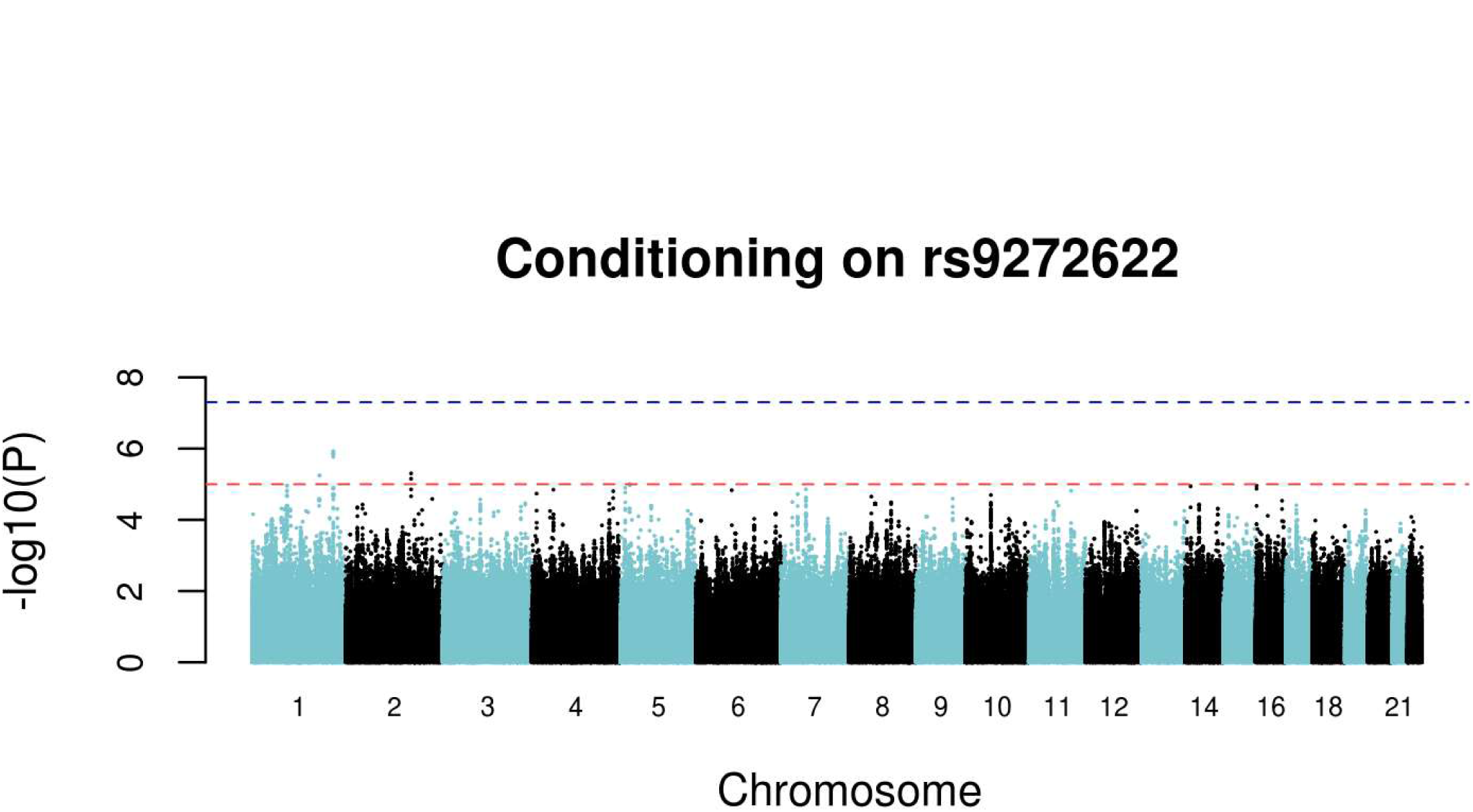
Manhattan plots of GWAS results for the 4.46M high quality 1000G imputed SNP variants after conditioning on the top SNP rs9272622. Data are for analysis in FastLMM looking for association between SNPs and RHD.

**Figure S3.**
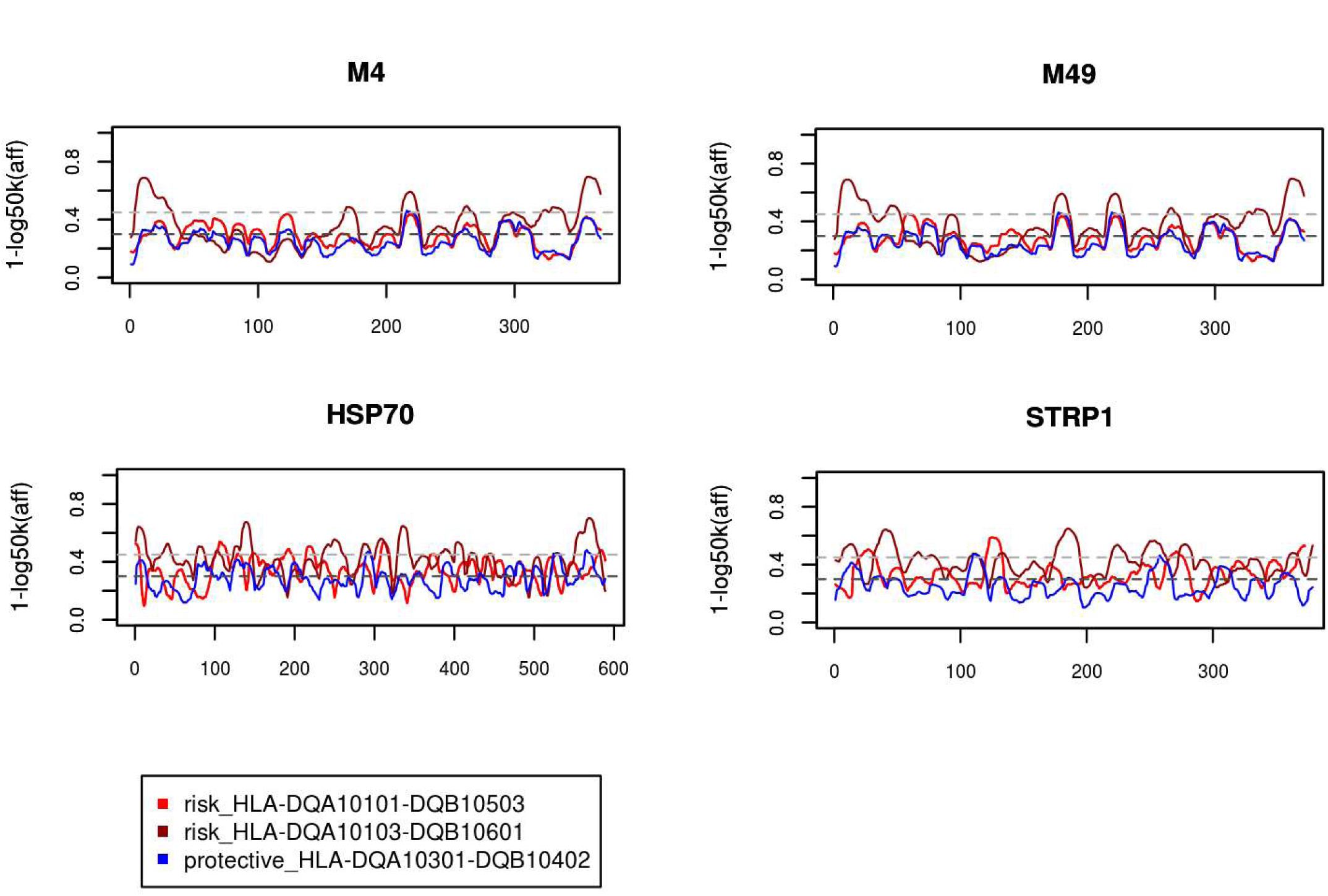
Plots showing binding affinities for predicted epitopes of GAS M proteins recognised by HLA DQ-DB heterodimers. Epitope binding predictions were performed in NetMHCIIPan3.1. The y-axis shows the relative binding affinity (expressed as 1-log_50,000_ of the nM binding affinity) for heterodimers formed from risk (red, brown) and protective (blue) DQ_DB haplotypes (see key); the x-axis indicates the amino acid sequence locations for mature proteins, also equivalent to the start position of overlapping 20mers (1-mer sliding window) in non-rheumatogenic GAS M4 (Accession number CAA33269) and M49 (Accession number AAA26868.1) sequences, GAS HSP70 (Accession number AAB39223.1) and GAS STRP1 (Accession number AAA26987.1) sequences. Horizontal lines indicate 500nM (upper) and 1000nM (lower) binding affinities.

**Table S1.**
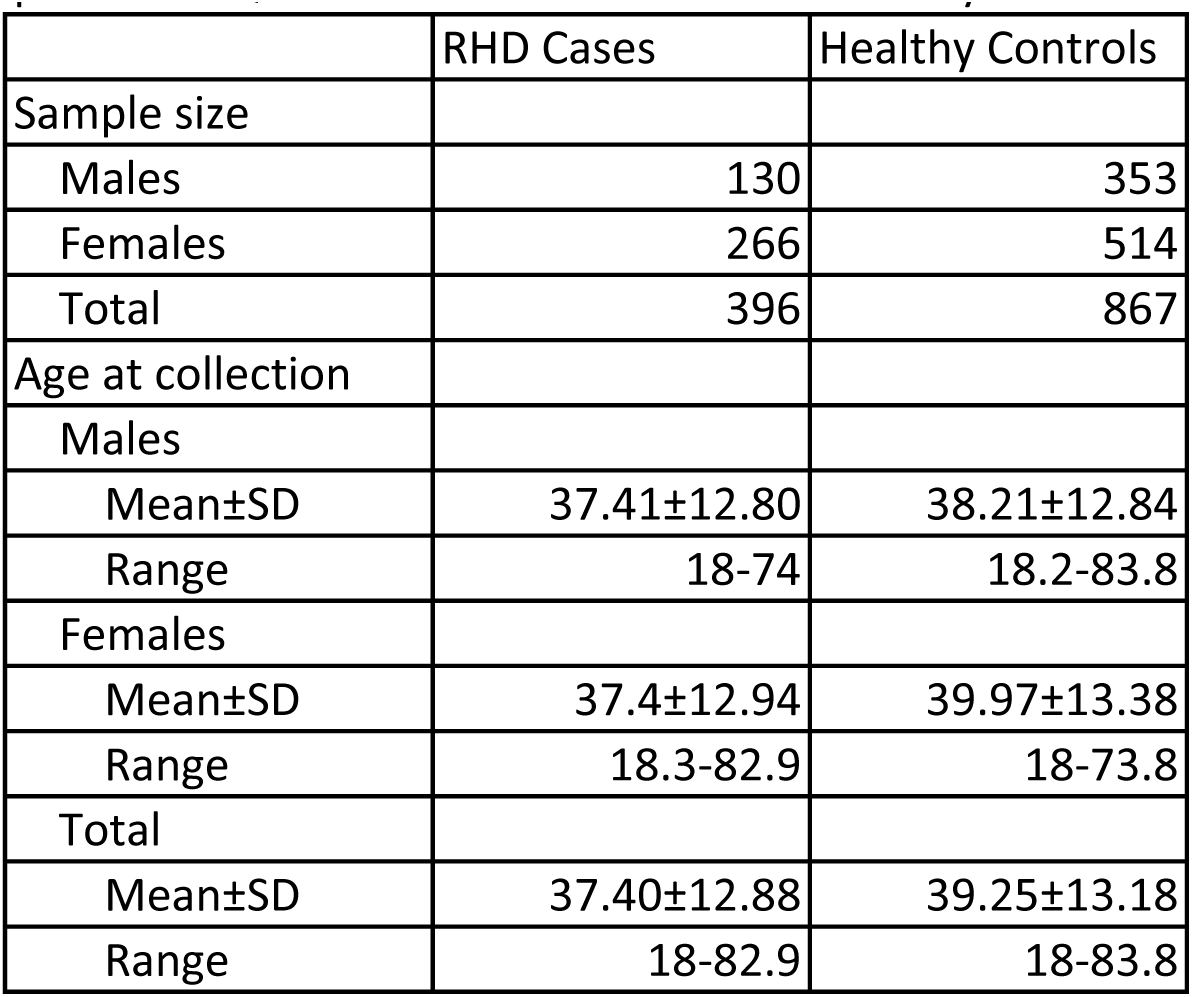
Basic demographic details (by gender, age at collection) for the 396 cases and 867 controls that passed all QC and were used in the GWAS analysis.

|  | RHD Cases | Healthy Controls |
| --- | --- | --- |
| Sample size |  |  |
| Males | 130 | 353 |
| Females | 266 | 514 |
| Total | 396 | 867 |
| Age at collection |  |  |
| Males |  |  |
| Mean±SD | 37.41±12.80 | 38.21±12.84 |
| Range | 18-74 | 18.2-83.8 |
| Females |  |  |
| Mean±SD | 37.4±12.94 | 39.97±13.38 |
| Range | 18.3-82.9 | 18-73.8 |
| Total |  |  |
| Mean±SD | 37.40±12.88 | 39.25±13.18 |
| Range | 18-82.9 | 18-83.8 |

**Table S2.**
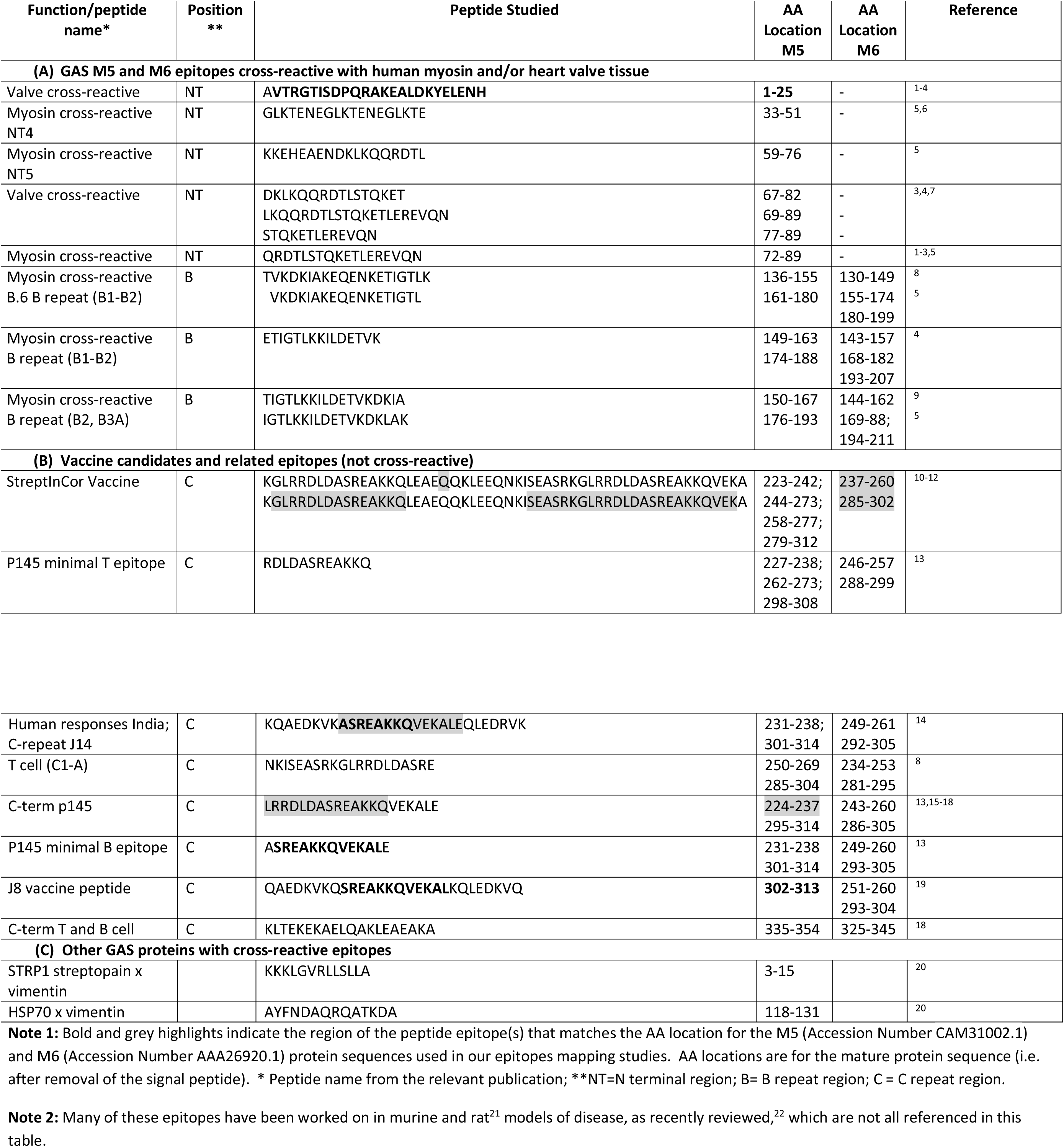
Summary of experimentally confirmed published epitopes for (A) GAS M proteins for which there is evidence of cross-reaction with human heart-related proteins. (B) GAS M proteins for which there is evidence of non-cross-reactive T-and B-epitopes, and (C) other GAS proteins with evidence of cross-reactive epitopes.

